# Acrylamide Induces Protein Aggregation in CNS through Suppression of FoxO1-mediated Autophagy

**DOI:** 10.1101/2022.07.29.498756

**Authors:** Xi-Biao He, Haozhi Huang, Yi Wu, Fang Guo

**Affiliations:** Laboratory of Stem Cell Biology and Epigenetics, School of Basic Medical Sciences, Shanghai University of Medicine and Health Sciences, Shanghai 201318, China.; Department of Orthopedic Surgery, Shanghai Tenth People’s Hospital Affiliated to Tongji University, Shanghai 200072, China.

## Abstract

Proteotoxic stress is a major stimulus and risk factor for the neuropathogenesis in central nervous system (CNS), which is tightly associated with neurodegenerative diseases. Here, we identify acrylamide (ACR), a type-2 alkene that is commonly detected in deep-fried starch food and used in water industry and biomedical laboratories, as a potent and universal inducer of proteotoxic stress in mouse brain, as well as in cultured mouse neural stem cells, neurons and astrocytes, three major cell types in CNS. Aggregations of ubiquitin-labeled misfolded proteins and neurodegeneration-related proteins including amyloid precursor protein and presenilin 1 were drastically induced by ACR, leading to cell-defensive aggresome formation that was able to temporarily counteract with ACR toxicity. However, a defected clearance of aggresomes eventually led to apoptotic cell death, which was largely attributed to the breakdown of cytoskeleton and impairment of macroautophagy/autophagy, as evidenced by an aggregation of filament actin and an almost complete loss of LC3-positive autophagosomes. A series of core autophagy-related genes responsible for autophagosome formation were down-regulated, indicating an ACR-induced transcriptional suppression of autophagy. In addition, FoxO1, the master transactivator of these genes, were both transcriptionally repressed and nuclear excluded by ACR. Overexpression of FoxO1 rescued ACR-induced autophagy defects and attenuated the proteotoxicity. In summary, we spotlight proteotoxic stress as a novel feature of ACR neurotoxicity in CNS, and implicate FoxO1 as a critical therapeutic target.

## INTRODUCTION

Proteotoxic stress-induced abnormal protein aggregation is a common cellular pathology in neurodegenerative diseases, which is caused by genetic deficits as well as environmental risk factors [45]. A balanced protein homeostasis requires an intact machinery for protein folding and misfolding, latter of which is largely attributed to protein degradation pathways including ubiquitin-proteasome system (UPS) and macroautophagy/autophagy [53]. Identifying novel proteotoxic stress risk factors that target these pathways paves avenues for the prevention and cure of proteotoxicity-related diseases.

Acrylamide (ACR), a type-2 alkene, is an environmental toxin to which the nervous system is especially vulnerable. In industry, ACR and its polymerization product polyacrylamide, are widely used as flocculant for clarification of potable water. Studies have showed that occupational workers exposed to acute ACR presented neurological manifestations in both peripheral nervous system (PNS) and central nervous system (CNS), including limb weakness, swelling, ataxia, confusion, disorientation, memory disturbance and hallucination [7]. To general public, the total exposure of ACR is significantly attributed to food and cigarette smoking, although the exposure rate is believed to be far from the threshold of acute exposure [39]. However, the accumulative feature of ACR still raises great concerns on the implication of chronic exposure of ACR in health and disease. Based on animal studies of ACR intoxication, the understanding of ACR neuropathology has evolved from distal axonopathy in PNS to neuronal nerve terminal degeneration and neurotransmitter inhibition in CNS [34]. A large number of in vitro studies have also confirmed the effect of ACR on the viability of neurons, neural stem cells (NSCs), astrocytes and microglia [14, 26, 32, 42, 55, 58], suggesting a general molecular mechanism of neurotoxicity might exist. Knowledge of organic chemistry has revealed that ACR rapidly forms irreversible covalent adducts at specific cysteine residues of proteins [36], suggesting that ACR might disturb normal protein homeostasis and functions. Moreover, several studies have reported aggregation of cytoskeleton components such as intermediate filament vimentin and neurofilaments induced by ACR [6, 40, 47, 48]. However, a direct link between ACR neurotoxicity and proteotoxic stress remains elusive.

Cells utilize a variety of molecular mechanisms to defend proteotoxic stress. When misfolded proteins aggregate, cells frequently assemble a vimentin-caged structure namely aggresome to temporarily wrap up protein aggregates for further degradation [19]. The assembly of aggresome is motorized by histone deacetylase HDAC6 and Dynein complex [21], whereas the disassembly and clearance is dependent on actin cytoskeleton remodeling [13]. The aggresomes and protein aggregates are subsequently submitted to and degraded through UPS and autophagy. Dysfunction of either step in this process leads to imbalanced proteostasis and impaired cell survival. For instance, vimentin-deficient NSCs presents proteasome mis-localization to aggresome, resulting in impaired proteostasis and delayed quiescence exit [37], whereas failed assembly of aggresome induced by HDAC6 deficiency worsens misfolded protein aggregation and apoptosis [21]. Blockade of filament actin (F-actin) remodeling, which is also regulated by HDAC6, exacerbates protein aggregation and neurodegeneration under proteotoxic stress, presumably due to failed fusion of autophagosome and lysosome for autophagic protein degradation [31]. In addition, disruption of F-actin dynamics evokes defensive cellular response of aggresome formation, impaired mitochondria-induced ROS production and eventually apoptotic cell death [29]. It remains unknown whether and how these mechanisms are involved in ACR neuropathology.

Regulation of autophagy occurs at both transcriptional and post-translational levels [28]. Repression of autophagy-related genes, by transcription factors (TFs) or histone modifiers, leads to failure in biogenesis and recycling of essential components for autophore or autophagosome formation [8]. Accumulating evidence have shown that many core autophagy-related genes are under transcriptional control of master TFs or histone modification, suggesting that these genes might share same upstream inducer and regulatory pathway. One of TFs mastering regulation of autophagy in this manner is Forkhead box O (FoxO) 1, member of FoxO family that are involved in a wide range of functions of cellular homeostasis and timing of aging [12]. FoxO1 is also a versatile player in CNS, whose roles include but not limited in maintenance and differentiation in NSCs [23, 41], prevention of axon degeneration and protein quality control in neurons [17, 54], inflammation modulation in astrocytes [52]. However, the association between environmental stress and FoxO1 dysfunction has not been well elucidated.

In this study, we have provided compelling evidence for ACR as a proteotoxic stress inducer in CNS, and identified FoxO1-mediated transcriptional regulation of autophagy as a key target of ACR. Our findings provide novel insights into ACR neurotoxicity and highlight proteotoxic stress as a potential risk of ACR exposure to CNS.

## RESULTS

### ACR induces protein aggregation in mouse brain

Intracellular proteostasis is critical for cell survival and injury defense. To establish a link between CNS proteostasis with neurotoxicity of ACR, we recruited a well-defined mouse model of acute ACR intoxication that applied intra-peritoneal injection of three doses of ACR for 5 days. The efficiency of this model was confirmed by an ACR dose-dependent decrease of motor coordination capacity in a rotarod test (Supplemental Fig. 1), which is in agreement with previous studies [25, 50]. To determine the level of misfolded proteins in different types of CNS cells in brain, ubiquitin was co-labelled with neuron-specific class III beta-tubulin (TuJ1) to mark neurons, with glial fibrillary acidic protein (GFAP) to mark astrocytes across several brain regions, and with Nestin to mark adult neural stem cells (NSCs) in the hippocampus (Fig. 1A). An ACR dose-dependent increase of ubiquitin expression in TuJ1+ neurons was observed in corresponding enriched regions including frontal association cortex, striatum, thalamus and cerebellum (Fig. 1B). Similarly, in the anterior and posterior corpus callosum, brain stem and cerebellum, where GFAP+ astrocytes were preferentially resided, ubiquitin expression was also specifically increased (Fig. 1C). In addition, ubiquitin expression was also induced in Nestin+ NSCs in the dentate gyrus (Fig. 1D). Importantly, most of ACR-induced ubiquitin were presented as dots rather than ubiquitous form in the cytosol, suggesting that they were protein aggregates. Consistent with these, Western blot analysis demonstrated that the total ubiquitin protein level in ACR-intoxicated mouse brains was significantly higher than that in control mice (Fig. 1E). These data indicate ACR as a potent and global inducer of misfolded proteins in CNS.

**Figure 1.**
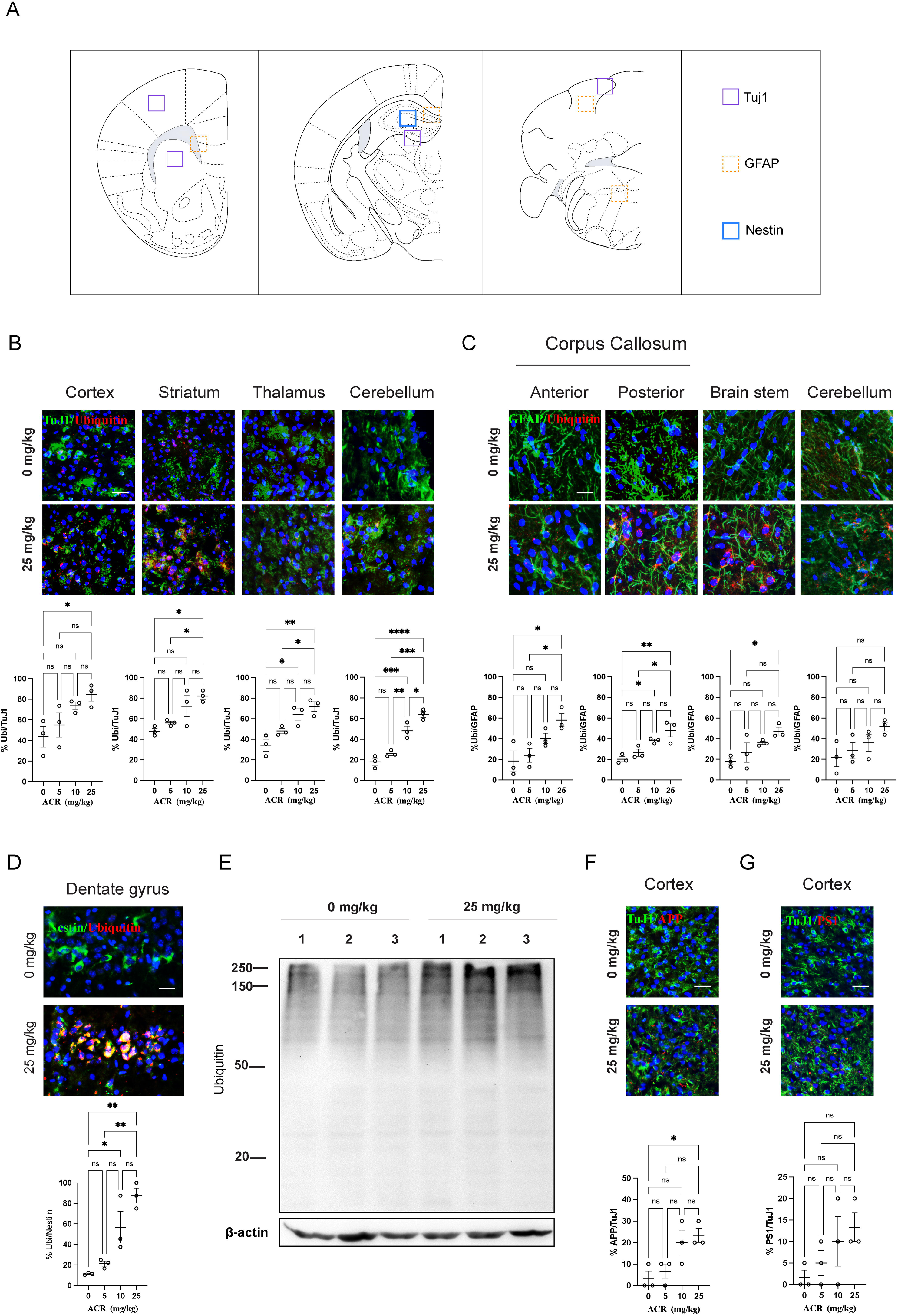
ACR intoxication induces global protein aggregation in mouse brain. (A) Schematics showing cell type-specific regions of adult mouse brain. (B) Representative immunofluorescence images and quantification of ubiquitin-expressing neuron-specific class III beta-tubulin (TuJ1)+ neurons in motor cortex, striatum, thalamus and cerebellum before and after 5 consecutive days of intraperitoneal ACR intoxication. (C) Representative immunofluorescence images and quantification of ubiquitin-expressing glial fibrillary acidic protein (GFAP) + astrocytes in anterior and posterior corpus collosum, brain stem, and cerebellum before and after ACR intoxication. (D) Representative immunofluorescence images and quantification of ubiquitin-expressing Nestin+ adult neural stem cells in dentate gyrus before and after ACR intoxication. (E) Western blot analysis of ubiquitin expression in whole brain extracts of mice with or without ACR intoxication. Each lane corresponds to one individual mouse brain. (F) Representative immunofluorescence images and quantification of amyloid precursor protein (APP)-expressing TuJ1+ neurons in motor cortex before and after ACR intoxication. (G) Representative immunofluorescence images and quantification of presenilin 1 (PS1)-expressing TuJ1+ neurons in motor cortex before and after ACR intoxication. Data represent mean ± S.E.M. **** P < 0.0001, *** P < 0.001, ** P < 0.01, * P < 0.05, ns, not significant; Two-way ANOVA with Tukey’s multiple comparisons test (n=3). Scale bar represents 20 μm.

Polyubiquitination of misfolded proteins precedes the degradation of many disease-associated proteins to maintain proteostasis in healthy brain. To assess the specificity of determine the effect of ACR on disease-associated proteins, TuJ1+ neurons in frontal association cortex were further co-labelled with amyloid precursor protein (APP) and presenilin-1 (PS1), two pathological proteins implicated in AD. Indeed, the cell number of TuJ1+ neurons expressing APP and PS1 were both largely increased after ACR intoxication in a dose-dependent manner (Fig. 1F, G). In addition, both proteins exhibited a form of aggregate. Collectively, our findings unravel an association of ACR intoxication with a global protein aggregation in CNS.

### ACR induces protein aggregation and apoptosis in cultured CNS cells

To confirm the effect of ACR on protein aggregation in vivo, we employed primary cell cultures of cortical NSCs, NSC-derived neurons and astrocytes (Supplemental Fig. 2) to investigate the expression and subcellular localization of ubiquitin-labeled proteins after exposure of ACR by immunoblotting and immunofluorescent analyses. In all three cell types, a time-dependent increase of ubiquitin expression was observed after cells were exposed to 10 mM ACR (Fig. 2A), and the ubiquitin was localized in the perinuclear regions of cells in the form of aggregates (Fig. 2B), suggesting the disrupted intracellular proteostasis as a common pathology for all three cell types in vitro. Notably, APP and PS1 were also aggregated in all three cell types after ACR treatment (Fig. 2C, D), suggesting a failed degradation of these pathological proteins. As a result, severe cell loss was found within 24 h of ACR exposure (Fig. 2E). DNA fragmentation marker TUNEL staining confirmed that ACR induced apoptosis in all three cell types (Fig. 2F). In addition, 7 days of sub-acute treatments of 0.05 mM ACR resembled the acute 10 mM concentration outcomes in astrocytes (Fig. 2G, H), indicating that the neurotoxicity of ACR in astrocytes, and likely in NSCs and neurons, was accumulative. Taken together, these results provide clear cell-based evidence of ACR as a strong inducer of proteotoxic stress in CNS.

**Figure 2.**
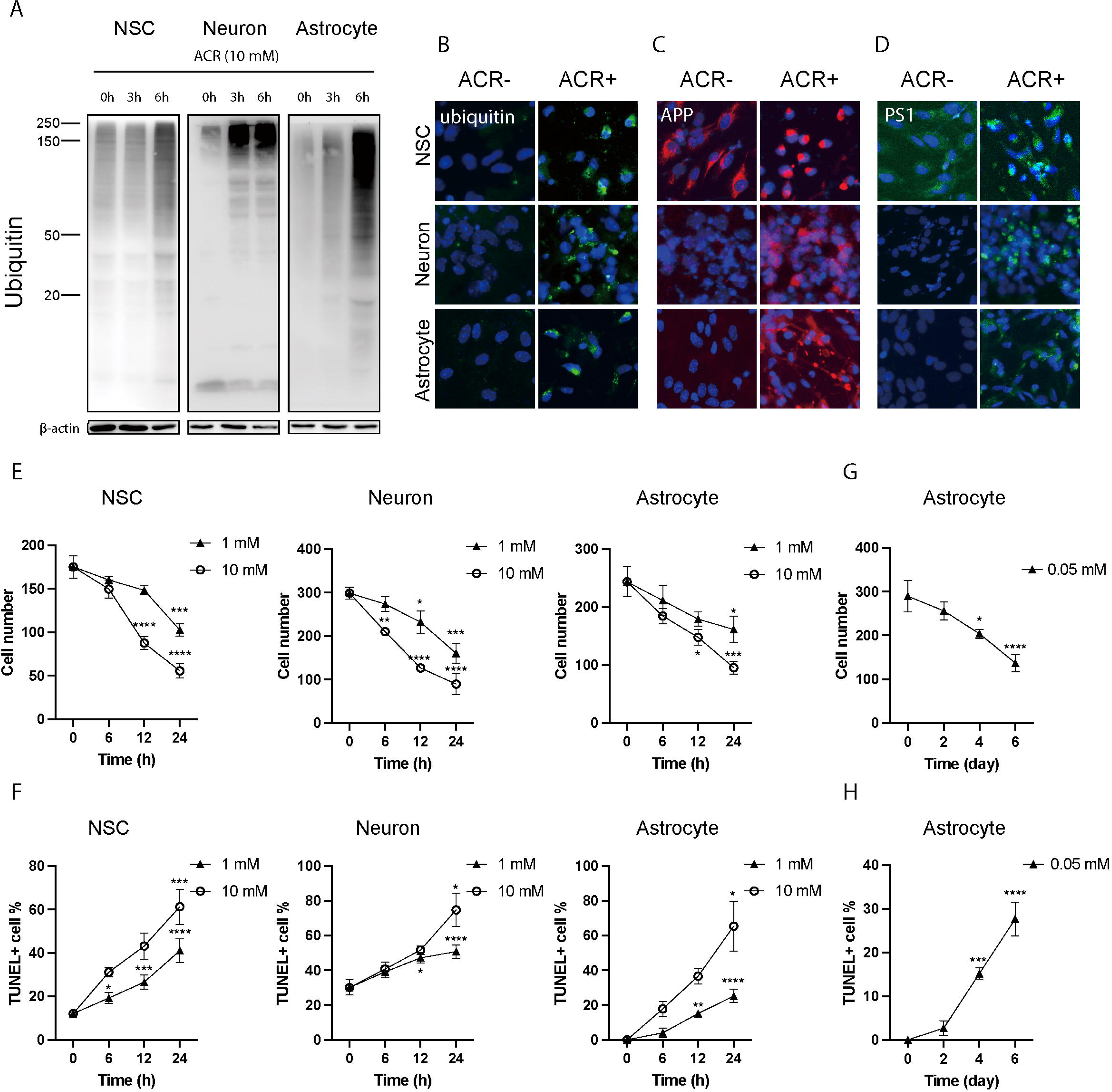
ACR proteotoxicity in vitro. (A) Western blot analysis of time-dependent ubiquitin expression in cultured primary neural stem cells (NSCs), neurons and astrocytes exposed to ACR. (B-D) Representative immunofluorescence images of ubiquitin, amyloid precursor protein (APP) and presenilin 1 (PS1) expression in cultured NSCs, neurons and astrocytes exposed to 10 mM ACR for 3 hours, respectively. (E, F) Quantification of viability and TUNEL-marked apoptosis of cultured NSCs, neurons and astrocytes exposed to two doses of ACR within 24 hours, respectively. (G, H) Quantification of viability and TUNEL-marked apoptosis of cultured astrocytes exposed to low-dose (0.05 mM) long-term (8 days) ACR, respectively. Data represent mean ± S.E.M. **** *P* < 0.0001, *** *P* < 0.001, ** *P* < 0.01, * *P* < 0.05, ns, not significant; Two-way ANOVA with Tukey’s multiple comparisons test (n=3). Scale bar represents 20 μm.

### Aggresome clearance is disrupted by ACR

We then sought to determine the molecular mechanism of ACR proteotoxicity. Aggresome formation and clearance is a general cellular defense mechanism evoked by misfolded protein stress that temporarily sequesters those proteins for further degradation [13, 21]. The process involves vimentin-caged aggresome assembly, microtubule-assisted aggresome migration to microtubule-organizing center (MTOC) region and actin-dependent fusion of autophagosome and lysosome at or in proximity to aggresomes. We first asked whether aggresome assembly and migration were functioning in response to ACR using cultured astrocytes as an example. Cells were visualized by co-labeling of ubiquitin and vimentin to determine whether ACR-induced protein aggregates were assembled into aggresomes. In normal condition, both proteins were ubiquitously present at basal level in the cytoplasm of astrocytes. From the early stage of ACR addition, small vimentin positive protein aggregates started to present in the perinuclear region, which overlapped with ubiquitin expression. The size of vimentin-positive aggregates gradually increased, forming large cytoplasmic foci, some of which were comparable to the size of cell nuclei (Fig. 3A). Vimentin structure positive for ubiquitin was similarly induced by ACR in NSCs and neurons (Suppl Fig. 3A). Further co-labeling analyses confirmed the composition of multiple disease-related proteins such as APP, PS1 and α-synuclein in those vimentin cages (Fig. 3B). Furthermore, they were aggregated at MTOC, evidenced by their co-localization with γ-tubulin. In addition, HDAC6 and Dynein, two molecules necessary for transporting misfolded proteins into aggresomes [21], were also co-localized with vimentin (Fig. 3C and Suppl Fig. 3B). These results confirmed ACR as a proteotoxic stress inducer and suggested an intact aggresome formation mechanism in response to ACR.

**Figure 3.**
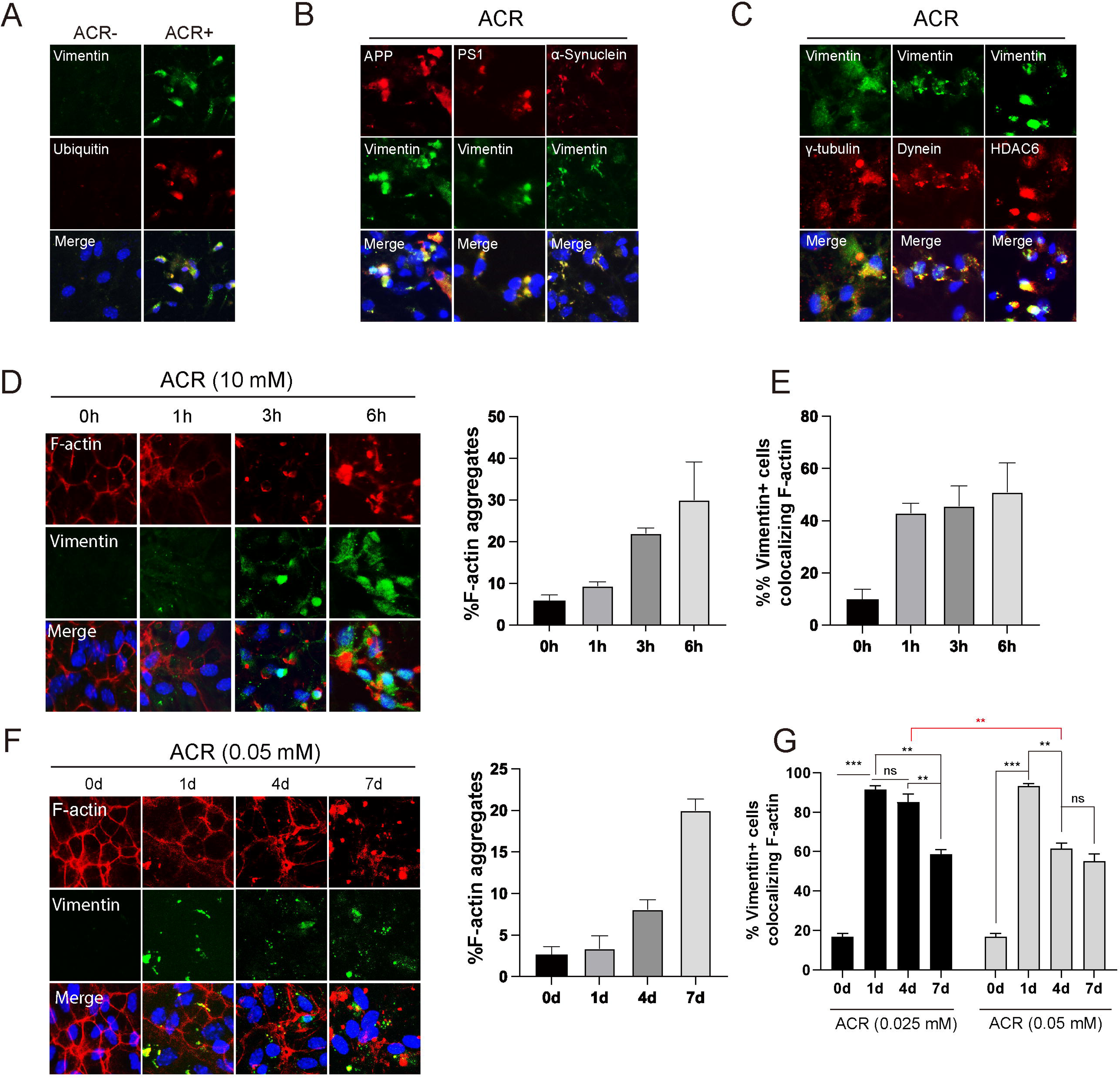
Defected aggresome clearance induced by ACR. (A) Representative immunofluorescence images showing co-localization of vimentin with ubiquitin in cultured astrocytes exposed to 10 mM ACR for 3 hours. (B) Representative images showing composition of disease-related protein amyloid precursor protein (APP), presenilin 1 (PS1), and α-synuclein in vimentin+ aggresomes in cultured astrocytes exposed to 10 mM ACR for 3 hours. (C) Representative images validating vimentin+ aggresomes localized at γ-tubulin+ microtubule organizing center and composed of aggresome motor complex components HDAC6 and Dynein after cultured astrocytes were exposed to 10 mM ACR for 3 hours. (D, E) Representative images and quantification of filament actin (F-actin) aggregates and their co-localization with vimentin in cultured astrocytes exposed to 10 mM ACR within 6 hours and exposed to 0.05 mM ACR within 7 days, respectively. Data represent mean ± S.E.M. *** *P* < 0.001, ** *P* < 0.01, ns, not significant; One-way ANOVA with Tukey’s post hoc test. Scale bar represents 20 μm.

A co-localization of F-actin network in aggresome is needed for subsequent autophagic clearance, as F-actin network has been reported to facilitate local autophagosome-lysosome fusion around aggresome [31]. To determine whether aggresome clearance mechanism was intact, we examined the F-actin dynamics in CNS cells in response to ACR by labeling F-actin with phalloidin. In normal condition, actin filaments are seen at the cell cortex and running linearly across the cell as stress fibers in astrocytes, NSCs and neurons. After ACR treatment, F-actin filaments were remarkably aggregated as puncta in astrocytes and NSCs, whereas their intensity in neurons were greatly decreased without forming aggregates, suggesting a severe actin cytoskeleton breakdown as a general CNS neuropathy in response to ACR (Fig. 3D and Suppl Fig. 3C). Notably, roughly 40% of vimentin-positive aggresomes co-localized with F-actin within 1 hour of 10 mM ACR treatment in astrocytes, suggesting an onset of aggresome-lysosome fusion. However, this rate was not further increased when examined at 3 and 6 hours of ACR exposure (Fig. 3E), despite that the size of ubiquitin-containing aggresomes continued increasing, suggesting a disrupted progress of aggresome-lysosome fusion. The loss of aggresome clearance capacity was more evident when astrocytes were exposed to long-term low dose ACR. As shown in Fig. 3E, the F-actin breakdown was not severe until the fourth day of 0.05 mM subacute ACR exposure. In consistent, the co-localization of vimentin+ aggresome with F-actin reached almost 100%, suggesting a complete function of aggresome-lysosome fusion. However, the co-localization rate was significantly decreased from the fourth day of ACR exposure. Reducing ACR concentration to 0.025 mM delayed the onset of this decrease (Fig. 3F), suggesting a positive correlation between ACR exposure and dysfunction of aggresome-lysosome fusion. These results collectively demonstrate an impaired molecular machinery of aggresome clearance related to autophagy.

### Autophagy is hampered by ACR

Aggresome clearance and disassembly is dependent on protein degradation pathways. Thus, we sought to determine whether UPS or autophagy-lysosomal pathway was impaired by ACR in astrocytes. UPS is the main protein degradation pathway critical for maintenance of intracellular protein balance. To determine the association of UPS and ACR-induced protein aggregation, proteasome inhibitor MG132 was treated to astrocytes exposed to ACR. The addition of MG132 markedly increased TUNEL labeling and ubiquitin expression in astrocytes (Suppl Fig. 4A), demonstrating a positive role of UPS in alleviating toxicity of ACR through degrading protein aggregates and reducing cell death. Moreover, a luminescent proteasome assay showed the proteasome activities represented by three major proteolytic activities in 20S core of proteasome including chymotrypsin-like, trypsin-like and caspase-like were either not altered or increased by ACR (Suppl Fig. 4B). These results suggest an active action of UPS in response to ACR.

**Figure 4.**
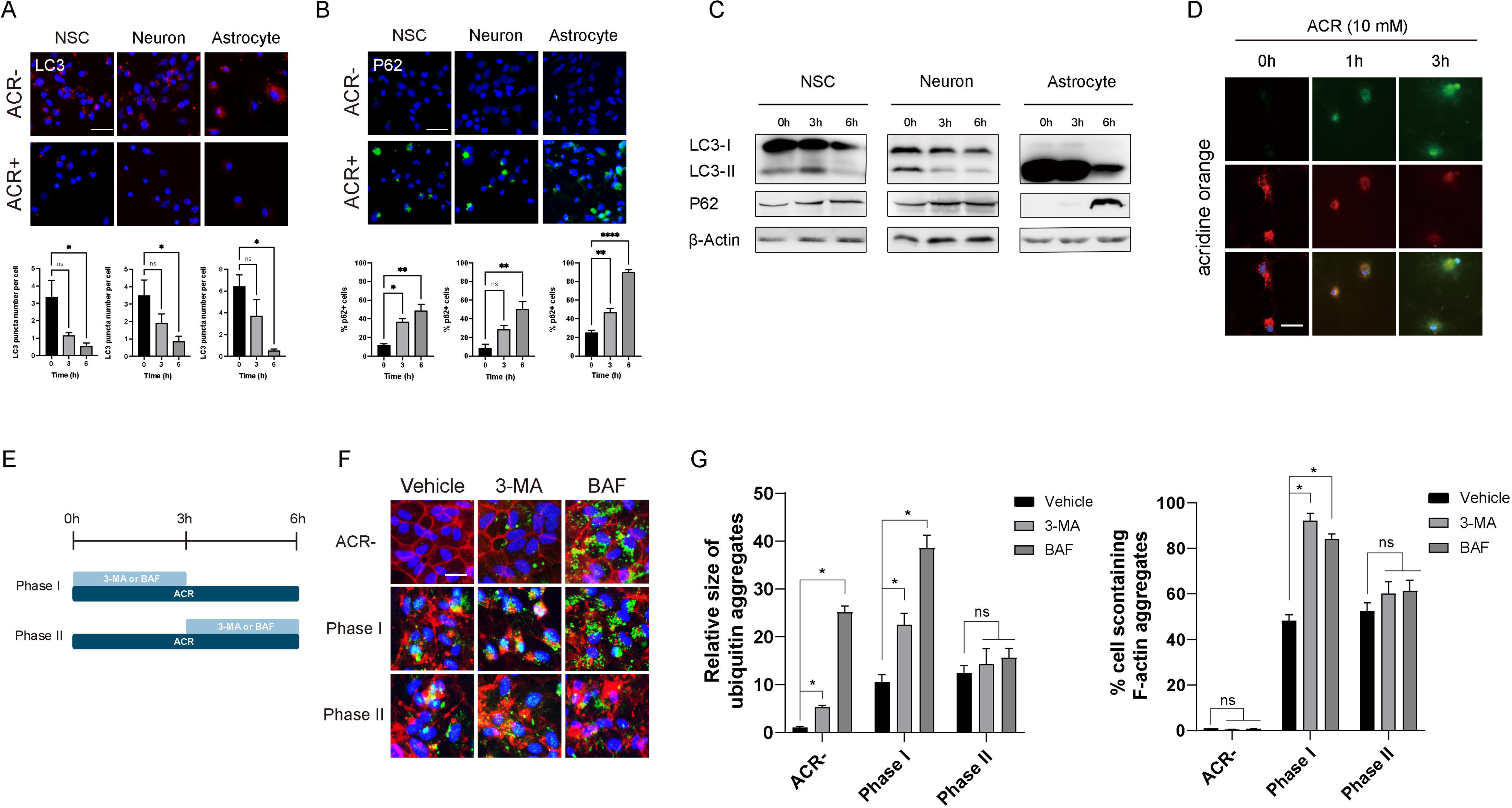
Disrupted autophagy induced by ACR. (A) Representative immunofluorescence images and quantification of LC3 puncta expression in cultured neural stem cells (NSCs), neurons and astrocytes before and after exposure to 10 mM ACR for 3 hours. (B) Representative images and quantification of p62 expression in three types of cells before and after same procedure of ACR exposure. (C) Western blot analysis of LC3 and p62 expression in three types of cells within 6 hours of ACR exposure. (D) Representative images showing astrocytes pre-treated with acridine orange and exposed to 10 mM ACR within 3 hours. Notice the gradual transition of cytosolic red fluorescence into nuclear and cytosolic green along ACR exposure, indicating the loss of acidic vesicles induced by ACR. (E) Schematics showing experimental procedure of stage-specific autophagic inhibition by treatment of 3-methyadenine (3-MA) or bafilomycin A (BAF) along with 6 hours of 10 mM ACR exposure in cultured astrocytes. (F) Representative images and quantification of ubiquitin expression and filament actin (F-actin) aggregation in astrocytes underwent experimental procedure above. Data represent mean ± S.E.M. **** *P* < 0.0001, ** *P* < 0.01, * *P* < 0.05, ns, not significant; One-way ANOVA with Tukey’s post hoc test. Scale bar represents 20 μm.

By contrast, we found a dramatic decrease of LC3 puncta in ACR-treated astrocytes, as well as in NSCs and neurons (Fig. 4A). The disappearance of LC3 puncta was indicative of a decrease of autophagy rather than an increased autophagic flux, as it was not recovered in the presence of bafilomycin A1 (BAF; data not shown). Consistently, autophagy cargo adaptor protein p62 which is also ubiquitin-binding protein, was markedly accumulated (Fig. 4B). Western blot analysis confirmed all the immunostaining observations above (Fig. 4C). Consequently, acidic lysosomes, the downstream executive organelles of autophagy exemplified in astrocytes, were disrupted, evidenced by the monitoring of acidic vesicles using acridine orange showing the dramatic loss of red lysosomes and increase of dispersed green in cytoplasm (Fig. 4D).

To search for direct evidence for the association between ACR toxicity and autophagy, we treated astrocytes with autophagy inhibitors 3-methyadenine (3-MA) or BAF. To determine the causal relationship between ACR and autophagy inhibition, inhibitors were treated in the early or late period of ACR exposure (schematic in Fig. 4E). Treatment of each inhibitor without ACR slightly increased ubiquitin expression without affecting the viability of cells, verifying a function of basal autophagy for normal protein turnover. As expected, inhibition of autophagy by either 3-MA or BAF in early period of ACR exposure (phase I) exaggerated ACR-induced aggregation of ubiquitin and F-actin, suggesting a positive role of autophagy in early-stage defense against ACR-induced proteotoxicity (Fig. 4F, G). However, the same thing did not happen when either of the inhibitors was treated in the late period of ACR exposure (phase II), when the progression of ubiquitin aggregation and F-actin breakdown already reached halfway. This result might reflect a progressive loss of autophagy upon ACR stimulation, given the diverse mechanisms of action of these drugs to inhibit autophagy. Nevertheless, these results demonstrate that ACR-induced proteotoxicity is largely associated with autophagy dysfunction.

### Transcriptional repression of autophagy-related genes by ACR

The almost complete loss of LC3-positive autophores and autophagosomes from immunostaining analysis suggested a deficit of their biogenesis. To determine the molecular mechanisms underlying this phenomenon, the transcriptional profile of genes related to autophagosome biogenesis [27] was monitored in cultured astrocytes upon ACR exposure. Dramatic down-regulations of genes responsible for autophagosome formation (pi3k, mtor, p70, beclin-1 and lc3b) and autophagosome maturation (atg5, atg7, atg9a, and atg12), were observed in a time-dependent manner (Fig. 5A). By contrast, genes responsible for autophagy initiation (ulk1, ulk2) and lysosome biosynthesis (lamp1, lamp2) were not altered (data not shown). Consistently, the majority of these alterations also occurred in vivo, evidenced by transcriptional analysis of mRNA extracted from motor cortex and brain stem regions of ACR-intoxicated or control mouse brains (Fig. 5B). These results indicated an association between ACR and transcriptional repression of autophagy.

**Figure 5.**
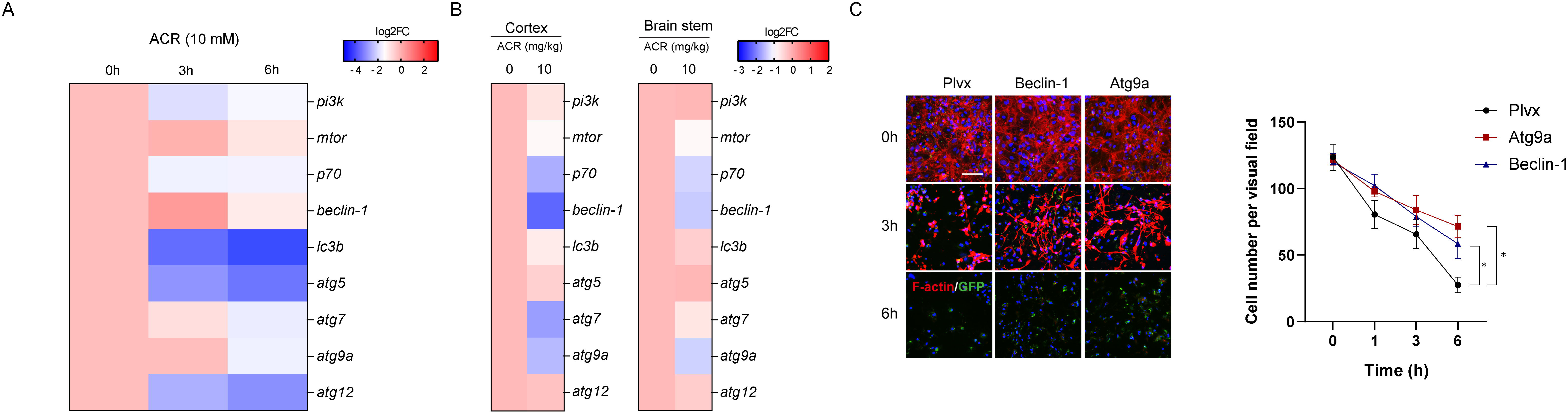
Transcriptional suppression of autophagy-related genes by ACR. (A) Heat map showing expression profile of core autophagy-related genes in cultured astrocytes exposed to 10 mM ACR within 6 hours. (B) Heat maps showing expression profile of core autophagy-related genes in two brain regions of mice underwent intraperitoneal injection of ACR. (C) Representative immunofluorescence images of filament actin (F-actin) and quantification of viability in astrocytes overexpressing Beclin-1, Atg9a and control Plvx exposed to 10 mM ACR within 6 hours. Cells carrying exogeneous gene were marked by GFP. Data represent mean ± S.E.M. * *P* < 0.05; One-way ANOVA with Tukey’s post hoc test. Scale bar represents 20 μm.

To further determine the role of autophagy-related genes in ACR toxicity, we selected beclin-1 and atg9a, two of the most down-regulated genes with diverse roles in autophagy, and performed gain-of-function experiments in ACR-treated astrocytes. Cells carrying GFP-tagged full-length mouse beclin-1 or atg9a demonstrated limited but statistically significant resistance to ACR toxicity and appeared to bear less severe F-actin breakdown in the early stage of ACR exposure (Fig. 5C). However, cells were eventually lost with almost complete breakdown of F-actin regardless of the exogeneous gene they carried. Importantly, the down-regulation of core autophagy-related genes induced by ACR was not altered by either overexpression of beclin-1 or atg9a (data not shown), suggesting that a regulation of a more upstream signaling that covered a wider range of autophagy machinery was required to achieve better rescuing outcome. Nevertheless, this result indicated that the transcriptional repression of autophagy-related genes was associated with ACR-induced toxicity.

### FoxO1 is the molecular target of ACR to repress autophagy-related genes

Multiple molecules have been identified as master regulators of autophagy at transcriptional level. Based on previous studies, FoxO family transcription factors are deeply involved in the transcriptional regulation of many core autophagy-related genes, which largely overlapped those repressed by ACR. Therefore, we asked whether the global repression of autophagy-related genes induced by ACR was resulted from regulation of FoxO levels. Indeed, FoxO1 transcription is dramatically down-regulated by ACR in astrocytes. By contrast, the transcription of another transcriptional regulator of autophagy TFEB remains unaltered. Importantly, the transcription of Bnip3, a mitophagy-related gene previously known to be a direct target of FoxO1, was also decreased (Fig. 6A). At the cellular level, FoxO1 protein was found normally localized in the nucleus of astrocytes. However, a significant cytosolic translocation and aggregation of FoxO1 was induced by ACR (Fig. 6B), suggesting that the subcellular localization of FoxO1 was regulated in response to ACR. On the other hand, the mitochondrion-bound Bnip3 was also found severely aggregated in the cytosol of ACR-treated cells, suggesting a mislocation and dysfunction of this protein and likely a failed clearance of impaired mitochondria in response to ACR (Fig. 6C)

**Figure 6.**
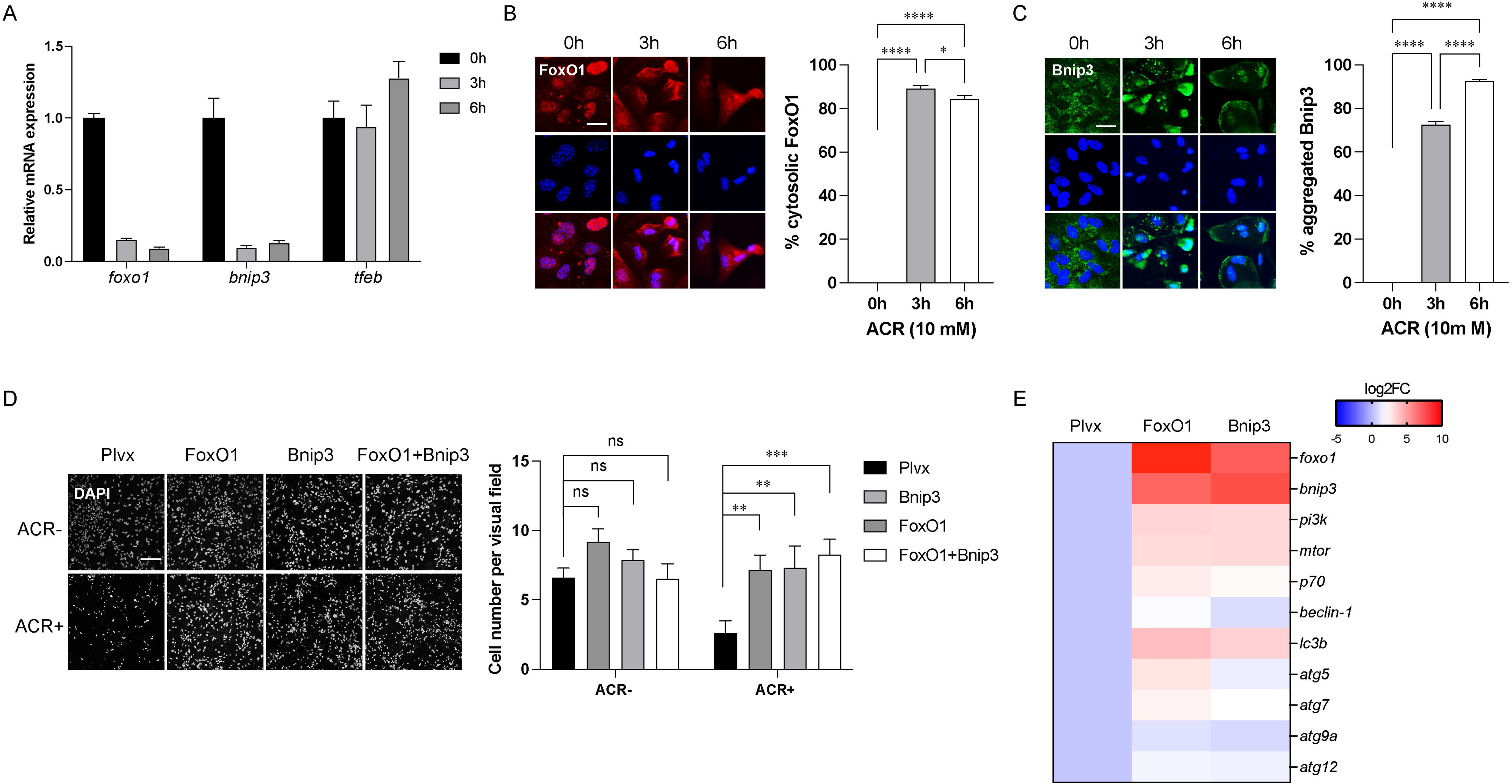
Repression of FoxO1 by ACR. (A) Real-time PCR analysis of expression change of foxo1, bnip3 and tfeb genes in cultured astrocytes exposed to 10 mM ACR within 6 hours. (B) Representative immunofluorescence images of FoxO1 expression and quantification of cytosolic FoxO1 in astrocytes exposed to 10 mM ACR within 6 hours. (C) Representative images of FoxO1 expression and quantification of aggregated Bnip3 in astrocytes exposed to 10 mM ACR within 6 hours. (D) Representative images of 4’,6-diamidino-2-phenylindole (DAPI)+ cell nuclei of astrocytes overexpressing FoxO1, Bnip3 or Plvx control exposed to 10 mM ACR for 6 hours. (E) Heat map of expression profile of autophagy-related genes in astrocytes overexpressing FoxO1, Bnip3 or Plvx control exposed to 10 mM ACR for 6 hours. Data represent mean ± S.E.M. N=3 independent experiments. **** *P* < 0.0001, *** *P* < 0.001, ** *P* < 0.01, * *P* < 0.05, ns, not significant; One-way ANOVA with Tukey’s post hoc test. Scale bar represents 20 μm.

To determine the role of FoxO1 and Bnip3 in ACR proteotoxicity, FoxO1 and Bnip3 were overexpressed through lentiviral delivery in astrocytes and challenged with ACR. Overexpression of either FoxO1 or Bnip3 greatly attenuated AR-induced cell death (Fig. 6D), indicating an anti-apoptotic feature of these two molecules. Furthermore, overexpression of both proteins altered the ACR-induced suppression of core autophagy-related genes such as atg5, atg7, beclin-1, atg12, and lc3b (Fig. 6E). Interestingly, overexpression of Bnip3 also up-regulated Foxo1 gene expression, suggesting that an interactive regulation of these two genes might be responsible to their similar effects in alleviating ACR proteotoxicity. Collectively, these results demonstrated that FoxO1 and Bnip3 was upstream factors of core autophagy-related genes that were transcriptionally targeted by ACR.

## DISCUSSION

The findings in this study demonstrate abnormal protein aggregation as a major CNS neuropathology of ACR exposure thereby identify ACR as a novel environmental risk factor of proteotoxic stress in CNS. ACR neurotoxicity is most commonly reported to affect the viability of CNS cells, including NSCs, neurons, astrocytes and microglia [4, 30, 33, 56, 59]. Besides, disturbance of astrocyte proliferation, NSC differentiation and neuronal neurotransmitter release have also been reported by subacute dose of ACR [15]. These effects might be attributed to direct or indirect outcomes from ACR-induced proteotoxic stress, given that the maintenance of intracellular protein homeostasis is critical for all the biological functions of cell. Importantly, several disease-specific proteins including APP, PS1 were severely accumulated after ACR exposure in vivo and in vitro. Although the dose of ACR applied to mouse in our experiments is several orders of magnitude higher than the estimated daily intake of ACR (0.3-0.8 μg per kg of body weight) for the general population, this finding still raise concerns on a potential association of accumulative subacute exposure with neurodegenerative diseases, particularly under conditions of low autophagy capacity such as metabolic disease and aging [24].

Chemical evidence have suggested that the neurotoxicity of acute ACR exposure is associated with its interaction with proteins, since it binds to sulfhydryl groups on cysteine residues of proteins forming irreversible adducts due to its high electrophilic property [35]. However, in case of subacute low-dose ACR exposure, one might argue that the defensive mechanisms of cell were potent enough to eliminate those protein adducts and prevent their aggregation. Indeed, aggresome formation is active in all three cell types and is able to temporarily resist against protein aggregation by sequestering small aggregates in vimentin-caged structure. However, our finding also reveals a transcriptional repression of autophagy underlying the failure of clearance of those aggregates, including gradually enlarged aggresomes. In comparison to the dynamic aggresome formation on protein level, this pathological mechanism on transcription level is more likely to be a sustained response to accumulative exposure of ACR and escalate the proteotoxicity regardless of ACR dose. This mechanism also suggests ACR as a long-term risk factor for inclusion body and plaque formation in the progress of neurodegeneration, which is highly associated with reduction of basal autophagy and gradual accumulation of pathological proteins.

Besides disease-specific proteins, ACR-induced protein aggregates are composed of various cytoskeletal proteins such as actin and intermediate filament vimentin. This finding is in consistent with previous studies reporting the stability and function of cytoskeleton as a main target of ACR in PNS [36, 47, 48]. The actin cytoskeleton provides mechanical support and movement force for cells, impairment of which usually leads to cell death [43]. Decrease of actin dynamics is also associated with depolarization of mitochondrial membrane and reactive oxygen species production and subsequent apoptosis [11]. In addition, actin cytoskeleton plays important roles in autophagosome formation and selective autophagy [1, 29, 31, 44]. Indeed, a direct association between ACR-induced actin aggregation and impaired fusion of aggresome and autophagy was observed in our work. It is reasonable to suspect that a wider range of actin-related biological functions including but not limited to those mentioned above are disturbed by ACR, emphasizing the importance of actin integrity for cell homeostasis. On the other hand, consistent with conclusions from previous studies [2, 20, 22, 38], we found that ACR-induced perinuclear aggregation of vimentin initially appeared as a defensive mechanism against misfolded proteins in the form of aggresome, which occurred in orchestration with an intact molecular machinery of aggresome formation including mobilization of HDAC6 and dynein and recruitment to MTOC. However, the size of aggresomes enlarged with ACR exposure, accompanied by failed fusion with autophagosomes, leading to exaggerated aberrant protein aggregation that greatly disturbed cell functions and survival. Our results have confirmed cytoskeleton as one of the major targets of ACR and added novel understanding of underlying molecular mechanisms for future therapeutic purposes.

To gain comprehensive insights into the neurotoxicity of ACR, we have validated most of the pathology in all three cell types in CNS. The similar response of these cell types to ACR despite of their diversity in molecular signatures and cellular functions suggest a common mechanism underlying ACR neurotoxicity. For technical convenience, mechanism explorations were performed on astrocyte, the cell type considered to have higher capacity to deal with stressed conditions than other CNS cell types [9, 10, 18]. Indeed, both in vivo and in vitro results suggest that astrocytes appear to exhibit stronger tolerance to ACR and higher basal autophagic activity than NSCs and neurons. In vivo, the ubiquitin accumulation in GFAP+ astrocytes in cortex and cerebellum is less induced by ACR than that in TuJ1+ neurons in same regions (Fig. 1A). In vitro, astrocytes exhibit lower rate of apoptosis in response to ACR (Fig. 2F) and a larger number of LC3 puncta under normal condition (Fig. 4A). In addition, astrocytes are able to dilute proteotoxicity through intercellular protein transfer [46, 51]. These facts might result in a higher vulnerability of NSCs and neurons to chronic ACR exposure, a neuropathology frequently found in neurodegenerative diseases. More evidence would be gained from future study comparing the effects of long-term low-dose ACR intoxication on different CNS cell types.

As the autophagy was inhibited in transcription level, it is likely that the conventional autophagy activator rapamycin, which works by inhibiting mTORC1, maybe not execute its proper function. Indeed, we found that pre-treating astrocytes with rapamycin only mildly delayed the onset of protein aggregation and apoptosis in the beginning of ACR exposure, but lost its effect after longer-time exposure (data not shown). Consistently, treating astrocytes with 3-MA or BAF in the later periods of ACR exposure did not worsen the toxicity. The best explanation for these observations is that the efficiency of those conventional autophagy regulation drugs is largely dependent on the level of basal autophagy and integrity of autophagy machinery, which remains active and intact in the beginning but completely lost in the end. Our results suggest that cautions should be taken when using drugs targeting post-translational regulation of autophagy for the treatment of conditions regarding transcriptional regulation of autophagy.

Our findings have established a link between the environmental proteotoxic stress with FoxO1 dysfunction in CNS. It remains unclear how ACR induces subcellular translocation and transcriptional repression of FoxO1. Stress stimuli such as oxidative stress usually trigger nuclear translocation of FoxOs, leading to transactivation of FoxO-mediated gene activation, which functions as adaptive or defensive mechanism providing beneficial effects to cells [3]. Therefore, the loss of nuclear and aggregation of cytosolic FoxO1 induced by ACR is likely to represent a pathological signature rather than an adaptive response, underscoring the importance of maintaining FoxO1 in nucleus against ACR proteotoxicity. The association of transcriptional repression of core autophagy genes with absence of nuclear FoxO1 confirms FoxO1 as a master transactivator of these genes in astrocytes and likely in other CNS cell types. On the other hand, the transcriptional repression of FoxO1 suggests an alternative mechanism underlying ACR-induced impairment of autophagy. Although the causal relationship between ACR-induced post-translational and transcriptional regulation of FoxO1 is unclear, it is well known that FoxO1 is widely involved in feedback regulatory loop with several downstream signaling molecules and transcription factors such as AKT and SIRT1 [5, 57]. It is plausible to propose an autofeedback loop between nuclear exclusion and transcriptional suppression of FoxO1. In addition, our finding has also suggested a positive feedback loop between FoxO1 and Bnip3, which might explain the mechanism by which overexpression of Bnip3 alleviated ACR proteotoxicity.

## Acknowledgements

This work was supported by the National Natural Science Foundation of China (31701287 and 32100779). We thank Yong Zhen and Yongjun Ma for technical help on data analysis and figure arrangement.

## Authors’ contributions

XBH conceived the study, performed the experiments and wrote the manuscript. YW and HH performed the experiments, collected and analyzed data, and performed figure arrangement. FG generated plasmids, collected and analyzed data. All authors have read and approved of the final version of the manuscript.

## Data Availability Statement

The data that support the findings of this study are available from the corresponding author upon reasonable request.

## MATERIALS AND METHODS

### Ethics

Adult ICR male mice (6 weeks) and neonatal ICR mouse pups were supplied by Shanghai Jiesijie Experimental Animal Co. All animal experiments were approved by the animal ethics committee of Shanghai University of Medicine & Health Sciences and have been performed in accordance with the ethical standards laid down in the 1964 Declaration of Helsinki and its later amendments.

### ACR intoxication and tissue preparation

ACR solution dissolved in distilled water was daily injected intraperitoneally into adult ICR mice at doses of 5, 10 and 25 mg/kg body weight for 5 continuous days. For RNA and protein extractions, mouse brains were extracted after cardiac perfusion with cold phosphate buffered saline (PBS) and homogenized by RNAiso Plus (TAKARA) and homemade RIPA buffer, respectively. For cryosection, mouse brains were fixed with 4% cold paraformaldehyde by cardiac perfusion and dehydrated with 30% sucrose. Tissues were then stored in -80L until they were cryosectioned into 14 μm thickness (Leica CM1950).

### Rotarod test

Mice after ACR intoxication were applied to an accelerating rotarod test according to manufacturer’s instruction (IITC). In detail, the 3 cm-diameter suspended rod was set to accelerate at a constant rate from 5 to 45 rpm in 240 s. Mice were pre-trained for 2 consecutive days before placed on the rod for a session of two repeats, with 180 s resting time allowed in between. A trial ended when the mouse fell off the rotarod or after reaching 240 s. The falling time was recorded and averaged of two repeats.

### Cell culture

Primary astrocytes were extracted from postnatal day 1 mouse cortices and cultured in Dulbecco’s modified Eagle medium supplemented with 10% fetal bovine serum and 1% Penecillin/Streptomycin in vitro, passaged once with a modified mild trypsin method [49] and underwent 2 days of trichostatin A treatment [16] to eliminate all the microglia contamination. Primary NSCs were extracted from embryonic day 13 mouse cortices, plated onto poly-L-ornithine and fibronectin double-coated culture materials and maintained in a modified N2 media supplemented with B27. Basic fibroblast growth factor and estrogen growth factor were daily added for proliferation (all culture media and supplements are from Thermo Fisher Scientific). Withdrawal of these factors led to spontaneous neuronal differentiation. Six days later, NSCs became neuronal cells. All cells were incubated in 5% CO2, 37℃ incubator. ACR solution ranging from 0.025 mM to 10 mM was treated for various time periods 1 hour after fresh media change to avoid discrepancy of autophagic activity.

### Plasmids and gene delivery

Mouse full-length Beclin-1, Atg9a, FoxO1 and Bnip3 have been inserted into plvx backbone vector to construct corresponding lentiviral vectors. The lentiviral vectors were introduced into H293T cells with packaging particles by transfection with Lipofectamine2000 (Invitrogen). Supernatant fractions were harvested 1 and 2 days after transfection, concentrated by PEG8000 and stored at -70°C until use. For viral transduction, cells were incubated with the viral supernatant containing polybrene (Sigma) for 6 hours (lentivirus), followed by a medium change.

### Reverse transcription and real-time PCR analysis

RNA preparation, cDNA synthesis and PCR analysis were performed as previously described [16]. The PCR primers are summarized in Supplementary Table S1.

### Immunofluorescence microscopy

Cells were fixed with 4% paraformaldehyde for 20 minutes. For immunostaining, fixed cells or tissue slices were permeabilized and blocked in PBS with 0.3% Triton-X100 and 1% bovine serum albumin for 40 minutes, incubated with primary antibodies diluted with blocking solution at 4L overnight. Alexa Fluor series of second antibodies (Thermo Fisher Scientific) were applied accordingly for one hour at room temperature. Cells were finally mounted in 4’,6-diamidino-2-phenylindole (DAPI) and examined using fluorescence microscope (Leica DMi8). The first antibodies are listed in Supplementary Table S2.

### Western blot analysis

Tissues were homogenized in home-made lysis buffer with protease inhibitor cocktails (Sigma). Cells were lysed for 30 minutes in same buffer. 3 μg of proteins from cell or tissue samples were separated by SDS-polyacrylamide gel electrophoresis and transferred to PVDF membrane. The membrane was blocked with 5% skim milk (Cell Signaling Technology) for 1 hour at room temperature, incubated with first antibodies including rabbit anti-ubiquitin, mouse anti-LC3, mouse anti-β-actin and rabbit anti-p62 (both from Cell Signaling Technology) overnight at 4 °C. The membranes were washed three times with Tris-buffered saline and Tween-20 followed by incubation with the peroxidase-conjugated anti-mouse or anti-rabbit secondary antibodies (both from Millipore) for 1 h at room temperature.

### Cell counting and statistics

For in vivo analysis, at least 50 immunoreactive or DAPI-stained cells were counted in each brain region. Data were derived from three mouse brains in same treatment group and expressed as mean ± S.E.M. For in vitro analysis, cells were counted in at least 10 random regions of each culture coverslip using an eyepiece grid at a magnification of 50 to 400X. Data are expressed as mean ± S.E.M. of three independent cultures. Statistical comparisons were made using Student’s t-test, one-way ANOVA with Tukey’s post hoc analysis or two-way ANOVA with Tukey’s multiple comparisons analysis (Graphpad Prism).

---

**Supplementary Figure 1. Rotarod test.** After five-day intraperitoneal intoxication of ACR ranging from 5 to 25 mg/kg body weight, mice were applied to an accelerating rotarod test to measure their motor coordination. n=5; data represent mean ± S.E.M. **** *P* < 0.0001, \**P* < 0.05, One-way ANOVA with Tukey’s post hoc test.

**Supplementary Figure 2. Cell culture validation.** Primary culture of neural stem cells (NSCs), NSC-derived neurons, and astrocytes are validated by immunofluorescence labeling with Nestin, TuJ1 and GFAP, respectively. Scale bar represents 20 μm.

**Supplementary Figure 3. Aggresome formation in neural stem cells (NSCs) and neurons.** (A) Representative immunofluorescence images of expression of ubiquitin and aggresome marker vimentin in NSCs and NSC-derived neurons exposed to 10 mM ACR for 3 hours. (B) Representative immunofluorescence images of expression of vimentin and HDAC6 and Dynein in NSCs and neurons exposed to 10 mM ACR for 3 hours. (C) Representative immunofluorescence images showing filament actin (F-actin) breakdown and aggregation in NSCs and neurons exposed to 10 mM ACR for 3 hours. Scale bar represents 20 μm.

**Supplementary Figure 4. Ubiquitin proteasome system (UPS) is not affected by ACR.** (A) Quantification of TUNEL+ cell percentage representing apoptosis in astrocytes in response to ACR and UPS inhibitor MG132. (B) Luminescent proteasome assay measuring three major proteolytic activities in 20S core of proteasome including chymotrypsin-like, trypsin-like and caspase-like in astrocytes exposed to 10 mM ACR within 24 hours. Data represent mean ± S.E.M. **** *P* < 0.0001, \**P* < 0.05, One-way ANOVA with Tukey’s post hoc test.

