## Supplementary figures and images for "Acrylamide Induces Protein Aggregation in CNS through Suppression of FoxO1-mediated Autophagy"

### Supplemental Figure 1

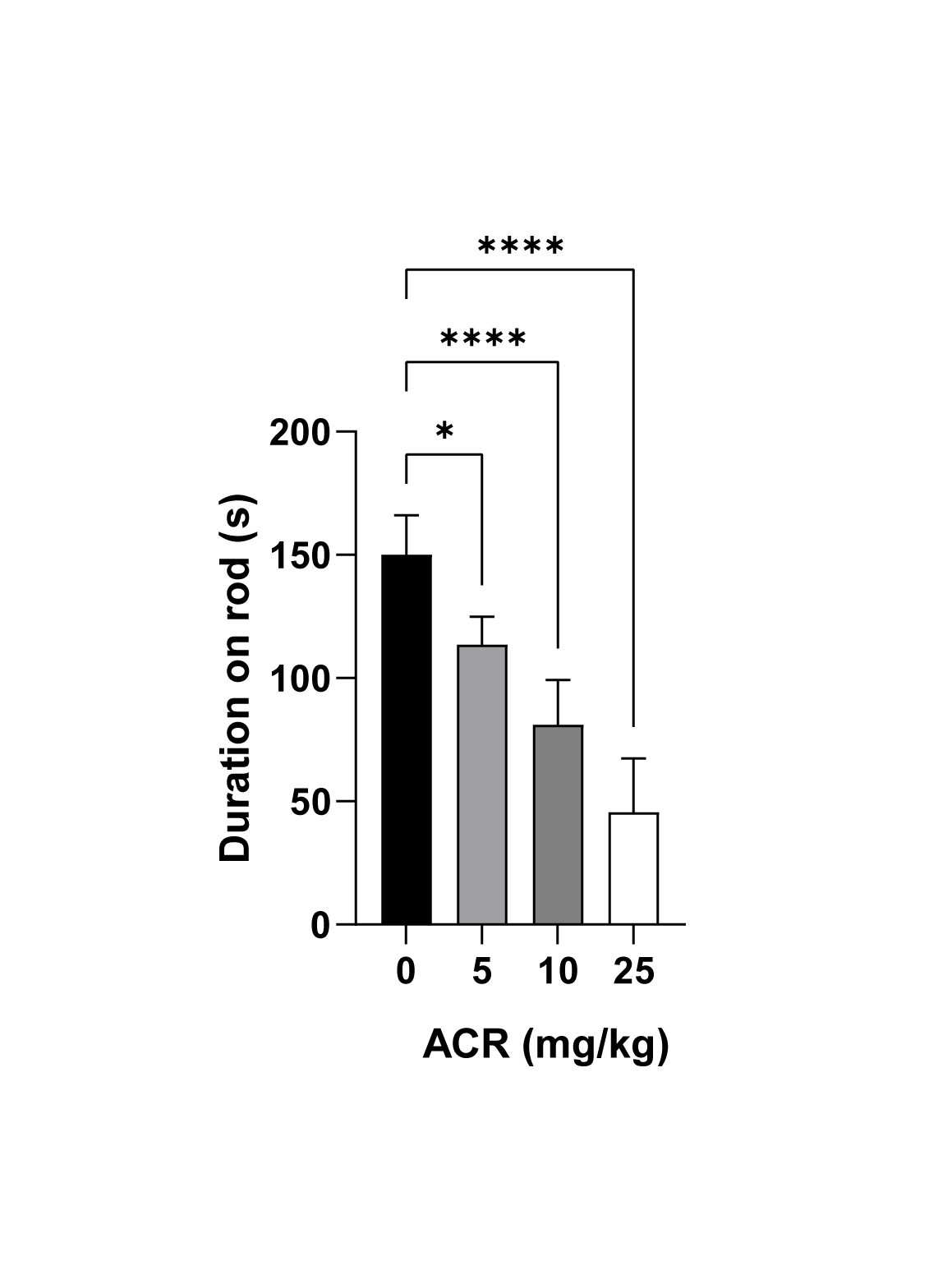

### Supplemental Figure 2

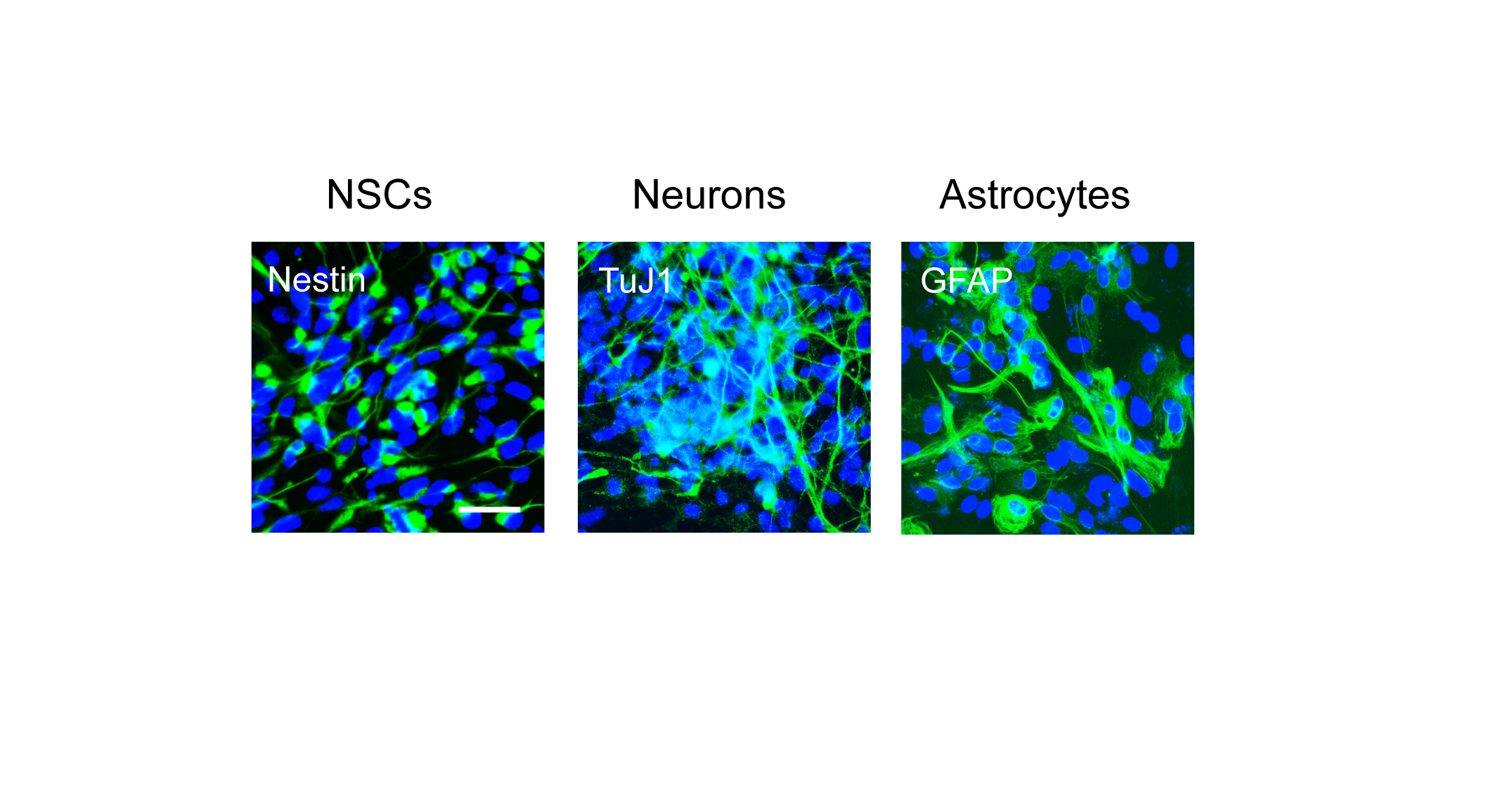

### Supplemental Figure 3

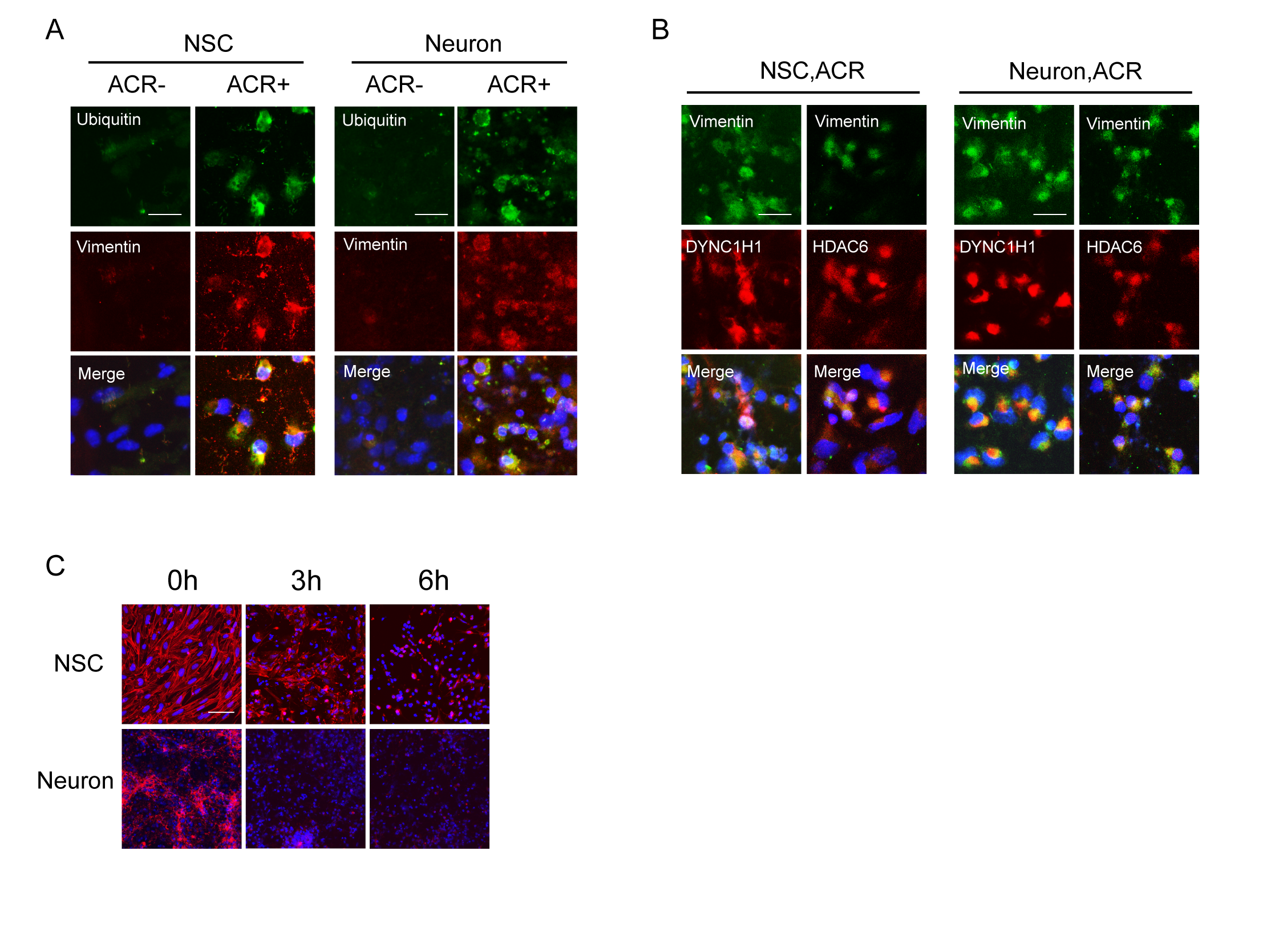

### Supplemental Figure 4

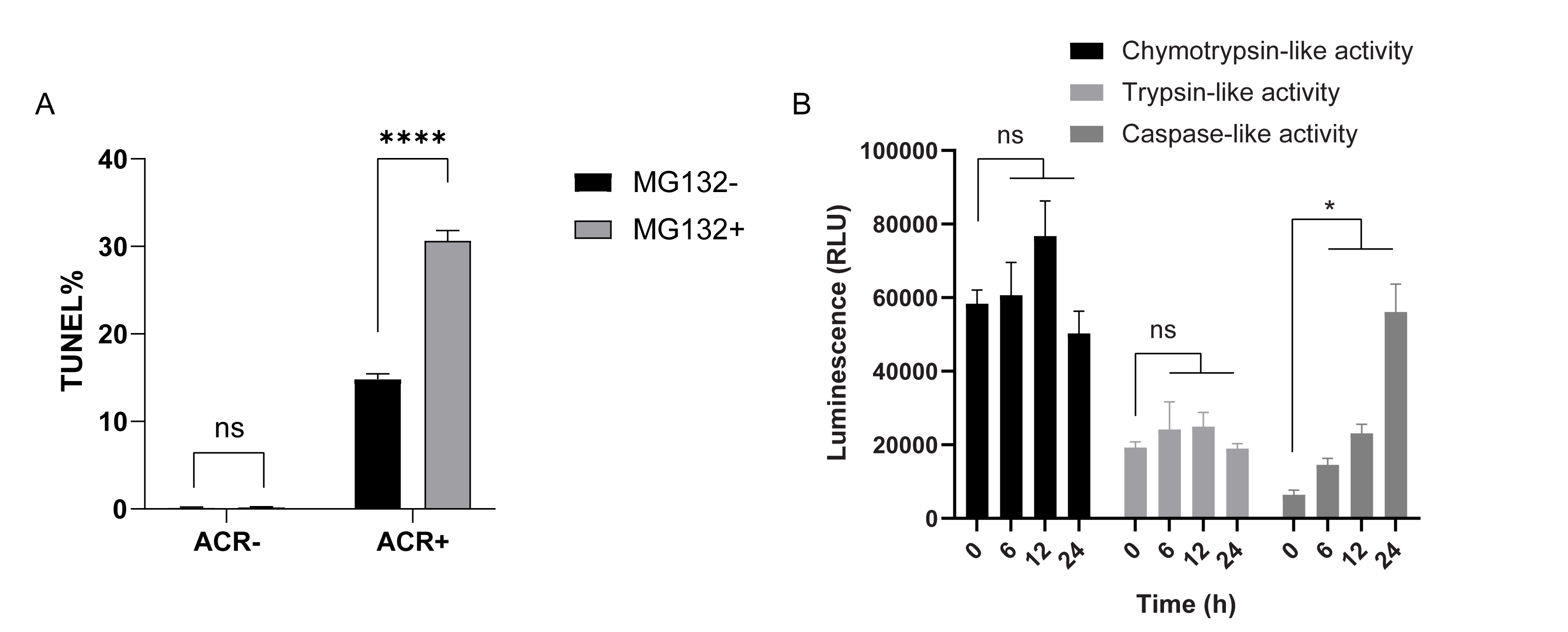
