## Supplemental Table 1 for "Acrylamide Induces Protein Aggregation in CNS through Suppression of FoxO1-mediated Autophagy"

**Supplemental Table S1.** Oligonucleotide primers

| Primer name | Forward | Reverse |
| --- | --- | --- |
| Atg5 | GCCCCTGAAGATGGAGAGAAG | GGTGGTTCCATCTAGCGAGG |
| Atg7 | GAGCGGCGGCTGGTAAGAA | TCTTCTGGGTCAGTTCGTGC |
| Atg9a | CCATCCTGGTCATTGCTGGT | ATGGAAGGGCAGACATTGGG |
| Atg12 | TGGCCTCGGAACAGTTGTTTA | ATCCCCATGCCTGGGATTTG |
| Pi3k | ACGTGCAGCTGAAGATAGGG | GGCAGGTCAGGGTACTTCAC |
| P70 | ACAGCGTGCTTTTACTTGGC | TGTGCGTGACTGTTCCATCA |
| Mtor | GGCCAAGCTACTGTGGCTAA | CAGATTGGATGGGTGCCTGT |
| Lc3b | AAGAGTGGAAGATGTCCGGC | TGCAAGCGCCGTCTGATTAT |
| Ulk1 | GCATCGAGCAAAACCTGCAA | GGGGAGAAGGTGTGTAGGGA |
| Ulk2 | AAGCGAGAGGCATCGATCTG | AGCATAACACCACAGGCTCC |
| Lamp1 | GCCCTGGAATTGCAGTTTGG | TGCTGAATGTGGGCACTAGG |
| Lamp2 | CTTAGCTTCTGGGATGCCCC | GCACTGCAGTCTTGAGCTGT |
| Bnip3 | CCCAGCATGAATCTGGACGA | TGAGAGTAGCTGTGCGCTTC |
| FoxO1 | CAGCAGCAACCCCTGTTTTC | CCCAGACAACTGCCCATGAT |
| Gapdh | TGTGAACGGATTTGGCCGTA | GTCTCGCTCCTGGAAGATGG |
| Beclin-1 | GGAAGTAGCTGAAGACCGGG | TTAGACCCCTCCATGCCTCA |
