## Supplemental Table 2 for "Acrylamide Induces Protein Aggregation in CNS through Suppression of FoxO1-mediated Autophagy"

**Supplemental Table S2.** Antibody list

| Primary antibodies | Supplier | Species | Reference |
| --- | --- | --- | --- |
| TuJ1 | R&D Systems | Mouse | MAB1195 |
| Ubiquitin | Dako | Rabbit | Z0458 |
| APP | Millipore | Mouse | mab348 |
| PS1 | Santa Cruz Biotechnology | Rabbit | 7860 |
| Gfap | Chemicon | Mouse | mab360 |
| Nestin | Millipore | Mouse | mab5326 |
| LC3 | MBL | Mouse | m152-3 |
| LC3 | MBL | Rabbit | pm036 |
| Phalloidin | Cell Signaling Technology |  | 12877 |
| β-actin | Cell Signaling Technology | Mouse | 3700 |
| P62 | Cell Signaling Technology | Rabbit | 5114S |
| Vimentin | Proteintech | Mouse | 10366-1-AP |
| α-Synuclein | Santa Cruz Biotechnology | Mouse | 12767 |
| γ-tubulin | Proteintech | Rabbit | 15176-1-AP |
| HDAC6 | Santa Cruz Biotechnology | Rabbit | 11420 |
| Dynein | Proteintech | Rabbit | 12345-1-AP |
